# Comparison of Approaches to the identification of Symptom Burden in Hemodialysis Patients Utilizing Electronic Health Records

**DOI:** 10.1101/458976

**Authors:** Lili Chan, Kelly Beers, Kinsuk Chauhan, Neha Debnath, Aparna Saha, Pattharawin Pattharanitima, Judy Cho, Peter Kotanko, Alex Federman, Steven Coca, Tielman Van Vleck, Girish N. Nadkarni

**Author notes:** TVV and GNN contributed equally. Lili Chan or Girish N. Nadkarni, MD, MPH Icahn School of Medicine at Mount Sinai, One Gustave L Levy Place, Box 1243, New York, NY 10029 Telephone number: Email Address or.

## Abstract

**Background:** Identification of symptoms is challenging with surveys, which are time-intensive and low-throughput. Natural language processing (NLP) could be utilized to identify symptoms from narrative documentation in the electronic health record (EHR).

**Methods:** We utilized NLP to parse notes for maintenance hemodialysis (HD) patients from two EHR databases (BioMe and MIMIC-III) to identify fatigue, nausea/vomiting, anxiety, depression, cramping, itching, and pain. We compared NLP performance with International Classification of Diseases (ICD) codes and validated the performance of both NLP and codes against manual chart review in a representative subset.

**Results:** We identified 1034 and 929 HD patients from BioMe and MIMIC-III respectively. The most frequently identified symptoms by NLP from both cohorts were fatigue, pain, and nausea and/or vomiting. NLP was significantly more sensitive than ICD codes for nearly all symptoms. In the Bio*Me* dataset, sensitivity for NLP ranged from 0.85-0.99 vs. 0.09-0.59 for ICD codes. In the MIMIC-III dataset, NLP sensitivity was 0.8-0.98 vs. 0.02-0.53 for ICD. ICD codes were significantly more specific for nausea and/or vomiting (NLP 0.57 vs. ICD 0.97, P=0.03) in BioMe and for depression (NLP 0.67 vs. ICD 0.99, P=0.002) in MIMIC-III. A majority of patients in both cohorts had ?4 symptoms. The more encounters available for a patient the more likely NLP was to identify a symptom.

**Conclusions:** NLP out performed ICD codes for identification of symptoms on several tests parameters including sensitivity for a majority of symptoms. NLP may be useful for the high-throughput identification of patient centered outcomes from EHR.

**Significance Statement:** Patients on maintenance hemodialysis experience a high frequency of symptoms. However, symptoms have been measured utilizing time-intensive surveys. This paper compares natural language processing (NLP) to administrative codes for the identification of seven key symptoms from two cohorts with electronic health records and validation through manual chart review. NLP identified high rates of symptoms; the most common were fatigue, pain, and nausea and/or vomiting. A majority of patients had ≥4 symptoms. NLP was significantly more sensitive at identifying symptoms compared to administrative codes for nearly all symptoms but specificity was not significantly different compared to codes. This paper demonstrates utility of a high throughput method of identifying symptoms from EHR which may advance the field of patient centered research in nephrology.

## Introduction

In the U.S. there are over 450,000 patients on hemodialysis.^1^ As mortality has decreased by over 30% over the past decade, improving the quality of life in HD patients has become a clinical and research priority. Symptom burden is extremely high in HD patients and from prior published survey data patients on average report a median of 9 symptoms over a seven day period.^2^ A recent initiative, the Standardized Outcomes in Nephrology (SONG)-HD, has identified outcomes important not only to physicians but also to patients.^3^ The top tier includes fatigue, cardiovascular disease, vascular access, and mortality. Middle and lower tier outcomes include symptoms such as pain, depression, anxiety, and cramps. While cardiovascular disease and mortality outcomes are easily tracked and identified, symptoms such as fatigue and depression are more difficult to identify and usually require prospective survey of patients. However, this method is low-throughput, time consuming (many surveys being over 30 questions), and only provides a view of the symptom burden at the time of survey administration.^4,5^

Electronic health records (EHRs) have been widely implemented in most major hospital systems and dialysis units.^6^ Granular clinical information for patient-centered care is routinely collected in EHRs. For example, during a hemodialysis treatment, patients are regularly observed for adverse signs and symptoms by nurses, technicians, and physicians. Additionally, they are under the care of a nutritionist and social worker. All of these encounters are routinely documented in EHRs as “free text” in progress notes and very infrequently as structured data such as International Classification of Diseases codes (ICD).^7^ While analysis of free text progress notes has traditionally been done via manual chart review, the advent of natural language processing (NLP) has the potential for the high throughput, rapid identification of symptoms from progress notes.

We undertook this study to determine the ability of NLP to retrospectively identify symptoms in HD patients from the EHR. We then compared the performance of NLP and ICD identification against manual chart review.

## Methods

### Study Population

From an original cohort of 38,575 participants from the Charles Bronfman Institute of Personalized Medicine Bio*Me* Biobank at the Icahn School of Medicine at Mount Sinai, we included patients with end stage renal disease (ESRD) who were on HD. The Bio*Me* Biobank is a prospective registry of racially and ethnically diverse patients that are recruited from primary care and subspecialty clinics in the Mount Sinai Healthcare System. The participants have provided their consent to have their EHR data available for biomedical research and linkage has been performed with the United States Renal Data System (USRDS) to ascertain dialysis status. The institutional review board approved the Bio*Me* protocols and informed consent was obtained for all subjects.

We retrieved all clinical notes of Bio*Me* Participants available from the centralized DataMart up to December 31, 2017. HD patients were identified as patients with ESRD according to the USRDS with exclusion of patients who received a kidney transplant and did not have a first dialysis date. Peritoneal (PD) patients were excluded using ICD 9 and 10 codes as PD and HD procedures are markedly different and associated with different symptoms and core outcomes **(Supplemental Table 1)**.

Additionally, we utilized the Medical Information Mart for Intensive Care (MIMIC-III) database to identify HD patients that were admitted to the intensive care unit. ^8^ MIMIC-III is a freely accessible critical care database of patients from a large, single center tertiary care hospital (Beth Israel Deaconess Medical Center in Boston, Massachusetts) from 2001 to 2012.^8^ This database includes patient demographics, billing codes, radiology reports, progress notes, and discharge summaries in deidentified form. We reviewed all clinical notes from the MIMIC–III database. As the MIMIC-III is a de-identified cohort, no linkage to USRDS could be done. Instead, ESRD was identified as patients who had an ESRD code and a code for dialysis procedure or diagnosis. PD patients were excluded by using PD procedure codes **(Supplemental Table 1).** As MIMIC-III is a de-identified publically available database, evaluation of data from this source was considered IRB exempt.

### Study Design

We utilized the CLiX NLP engine produced by Clinithink (London, UK) to parse documents from HD patients in the Bio*Me* Biobank and in MIMIC-III. There was no restriction on number of notes or types of notes placed. CLiX NLP is a NLP software that parses through free text and matches it to SNOMED clinical terms.^9^ SNOMED is a comprehensive healthcare terminology resource that is used in over fifty countries around the world. SNOMED has an inherent hierarchy consisting of overarching concepts, i.e. parent terms, which encompass more specific concepts, i.e., children terms. **Supplemental Figure 1** includes an example of how “cramp” would be represented in the SNOMED hierarchy and a search for cramps would also identify the seven children terms of bathing cramp, cramp in limb, cramp in lower limb, hand cramps, heat cramps, recumbency cramps, and stomach cramps. CLiX NLP is also equipped to handle typographical errors, sentence context, and negation.

We selected clinical outcomes from all tiers of outcomes identified from the SONG-HD initiative.^3^ Specifically, we queried for the following: fatigue, depression, pain, nausea and/or vomiting, anxiety, and cramps. The list of SNOMED concepts and children terms is included in **Supplemental Table 2**. These specific terms were selected due to their inability to be identified from structured data as opposed to outcomes such as hospitalizations, mortality, and dialysis adequacy, which can be identified using administrative codes or other structured data.

CLiX NLP mapped text from clinical notes to the corresponding SNOMED clinical terms. This was first identified on the document level and then on the patient level. For fatigue, pain, nausea and/or vomiting, anxiety, and cramps, NLP identification in at least one note was considered as test positive. For depression, as the disease is more chronic in nature, NLP identification in at least two notes on at least two different dates was necessarily to be considered test positive. We performed two iterations of NLP parsing with manual chart review guiding the second iteration. We rectified errors in identification in the NLP engine prior to the execution of the final parsing. Examples included phrases such as “The patient was advised to call for any fever or for prolonged or severe pain or bleeding” and “EKG sinus tach with V4, V5 depressions”. We modified the NLP algorithm to recognize these as negative expressions. We report results in this manuscript from the final NLP query.

We compared performance of ICD-CM codes with the results obtained from CLiX NLP. ICD-CM codes were chosen as described in prior literature when available and through physician extensive review of ICD-CM codes when not available. ^10–12^ ICD-9 and 10 codes were used in BioMe while only ICD-9 codes were available in MIMIC-III **(Supplemental Table 3).** Finally, both methods were compared with independent chart review by two physicians (LC and KB). When there was disagreement between manual validations for a patient, joint review of the patient’s chart was performed until consensus agreement was obtained.

As an additional validation of our NLP results, specifically for the depression query, we identified patients who had undergone Patient Health Questionnaire 9 (PHQ-9) screening since this is a common survey based instrument that is administered to dialysis patients.^13,14^ We considered depression screening positive if patients scored ≥10. If scores were <10 or there was discrepancy between depression as identified by PHQ-9, ICD, or NLP, additional manual chart review was done to identify evidence of history of depression (i.e. cognitive behavior therapy, anti-depressive medications, or prior suicide attempts). For patients with multiple PHQ9 scores, the highest score was used.

### Statistical Analyses

We calculated sensitivity, specificity, positive predictive value (PPV), negative predictive value (NPV), and F1-score of NLP and ICD9/10 codes. For cells on the 2×2 table where the value was 0, we entered 0.5 to allow for calculation of test statistics. We compared estimates of sensitivity, specificity, and F1-score using the McNemar test with significance set using a two sided p value of <0.05. We compared NPV and PPV using the generalized score statistic.^15^ All analyses were done using SAS version 9.4 (SAS Institute, Cary NC).

## Results

### Patient Characteristics

Out of 1152 patients with ESRD identified by the USRDS in Bio*Me*, we identified 1034 (90%) patients receiving maintenance HD. These HD patients had a mean age of 63.6±13.3 years, 42% were women, and 42% self-reported as African American. As expected of HD patients, there was high prevalence of diabetes (51%), hypertension (85%), coronary artery disease (53%), and congestive heart failure (31%) **(Table 1).** The median number of encounters was 109, (interquartile range [IQR] 41-241).

**Table 1:** Patient Characteristics of BioMe and MIMIC-III

|  | **BioMe (n=1034)** | **MIMIC-III (n=929)** |
| --- | --- | --- |
| Age [years] | 63.6 ±13.3 | 67.4±37 |
| Female | 433 (42) | 384 (41) |
| Race/ethnicity: |  |  |
| African American | 433(42) | 207 (22) |
| European American | 146(14) | 581 (63) |
| East Asian | 14(1.4) | 30 (3) |
| Hispanic | 376(36) | 40 (4) |
| Missing | 2(0.2) | 46 (5) |
| Other | 63(6) | 25 (3) |
| Comorbidities: |  |  |
| Diabetes | 526(51) | 544 (59) |
| Hypertension | 881(85) | 855 (92) |
| Coronary artery disease | 550(53) | 459 (49) |
| Congestive heart failure | 327(31) | 530 (57) |

Data are shown as mean ± standard deviation or count (%)

From MIMIC-III, we identified 929 HD patients utilizing ICD-9 codes. The mean age of patients was 67.4±37 years, 41% of patients were women, and 63% self-reported as white. Prevalence rates of chronic co-morbidities were similarly high, diabetes (59%), hypertension (92%), coronary artery disease (49%), and congestive heart failure (57%) **(Table 1).** Encounter analysis could not be done in MIMIC-III as the database only included encounters with ICU stays and nearly 80% of patients had only one ICU admission.

### Symptom Identification using NLP vs Administrative Codes

In BioMe HD patients, NLP identified symptoms more frequently than did ICD-9 and 10 codes **(Figure 1A).** The most frequent symptoms identified were pain (NLP 93% vs. ICD 46%, P<0.001), fatigue (NLP 84% vs. ICD 41%, P<0.001), and nausea and/or vomiting (NLP 74% vs. ICD 19%, P<0.001). The symptoms that were picked up best by both NLP and ICD were pain (45%), fatigue (40%), and depression (19%).

**Figure 1:**
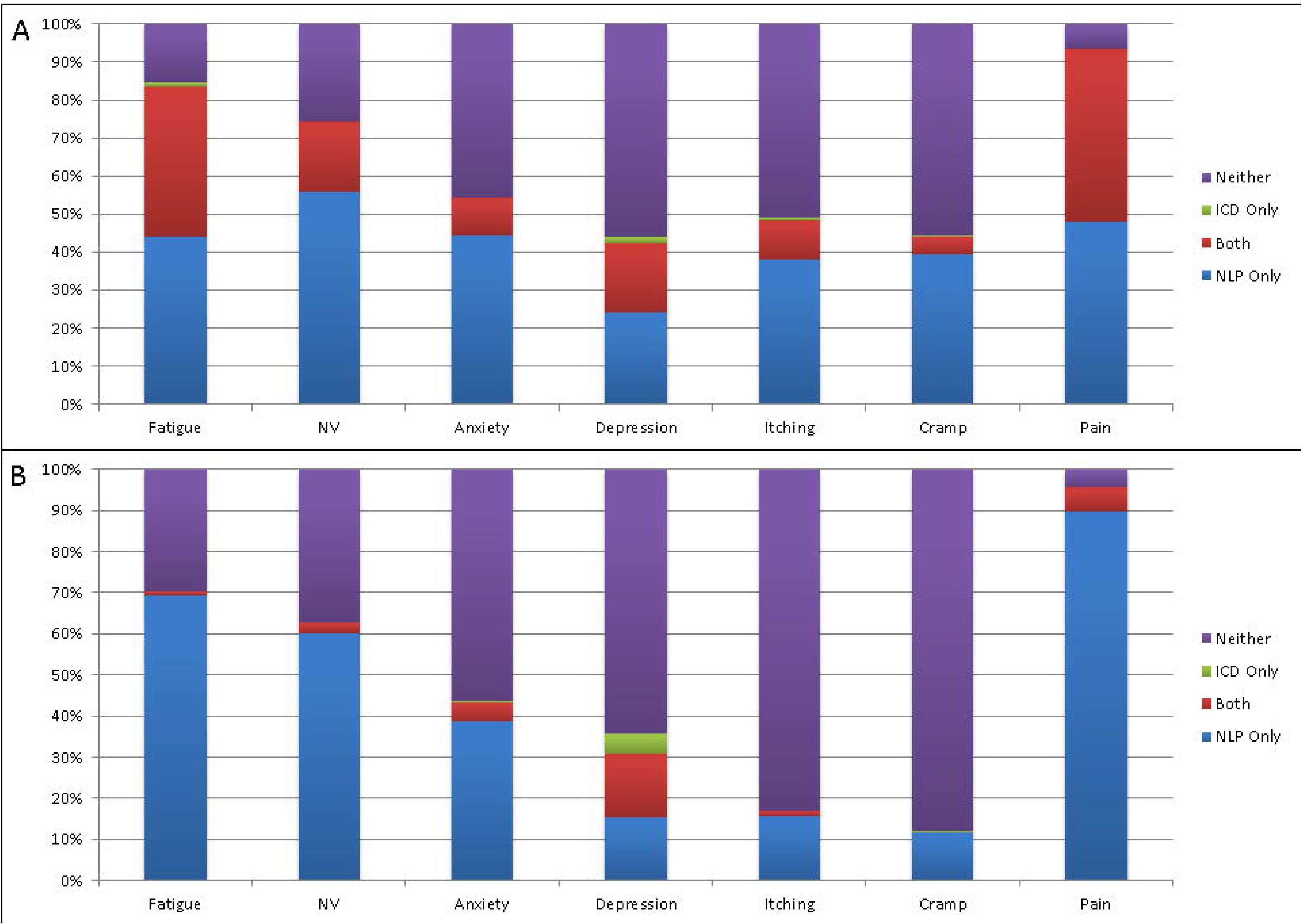
Frequency of symptom identified by NLP and ICD from (A) Bio*Me* and (B) MIMIC-III. Blue bar indicates percentage of patients where symptom was found only by NLP, green bar indicates percentage of patients where symptom was found by only by ICD, red bar indicates percentage of patients where symptom was found by both NLP and ICD, while purple bar indicates percentage of patients where the symptom was found by neither NLP or ICD.

In the MIMIC-III cohort, again NLP identified symptoms more commonly than ICD-9 codes. **(Figure 1B)** The symptoms with the highest prevalence according to NLP were pain (NLP 96% vs. ICD 6%, P=0.16), fatigue (NLP 70% vs. ICD 41%, P<0.001), and nausea and/or vomiting (NLP 63% vs. 19%, P <0.001). ICD-9 codes were best able to identify depression (17%) and no ICD-9 code for cramps was found in MIMIC-III.

### Manual Chart Validation

Overall, NLP was superior to ICD for identifying symptoms in both cohorts. In the Bio*Me* dataset sensitivity for NLP ranged from 0.85 to 0.99 while sensitivity for ICD ranged from 0.09 to 0.59 for ICD. In the MIMIC-III dataset, sensitivity for NLP ranged from 0.8 to 0.98 while sensitivity for ICD ranged from 0.02 to 0.53. **(Table 2 A/B)**

**Table 2:** Sensitivity, specificity, PPV, NPV, and F1 score of NLP vs. ICD for identification of symptoms for (A) BioMe and (B) MIMIC-II

|  | Fatigue |  |  | Nausea and /or vomiting |  |  | Anxiety |  |  | Depression |  |  |
| --- | --- | --- | --- | --- | --- | --- | --- | --- | --- | --- | --- | --- |
|  | NLP (95% CI) | ICD (95% CI) | P | NLP (95% CI) | ICD (95% CI) | P | NLP (95% CI) | ICD (95% CI) | P | NLP (95% CI) | ICD (95% CI) | P |
| **Sensitivity** | 0.99 (0.93-1) | 0.59 (0.43-0.73) | <0.001 | 0.99 (0.9 – 1) | 0.25 (0.12-0.42) | <0.001 | 0.98 (0.88 -1) | 0.3 (0.15-0.5) | <0.001 | 0.85 (0.65-96) | 0.54 (0.33-0.73) | 0.02 |
| **Specificity** | 0.89 (0.4-1) | 0.75 (0.19-1) | 0.68 | 0.57 (0.29-0.82) | 0.97 (0.77-1) | 0.03 | 0.9 (0.68-0.99) | 0.98 (0.83-1) | 0.22 | 0.81 (0.53-0.9) | 0.96 (0.79-1) | 0.06 |
| **PPV** | 0.99 (0.92-1) | 0.96 (0.82-1) | 0.3 | 0.86 (0.71-0.95) | 0.95 (0.66-1) | 0.02 | 0.94 (0.79-0.99) | 0.95 (0.66-1) | 0.16 | 0.79 (0.59-0.92) | 0.93 (0.68-1) | 0.12 |
| **NPV** | 0.89 (0.4-1) | 0.13 (0.03-0.35) | 0.007 | 0.94 (0.63-1) | 0.34 (0.2-0.51) | <0.001 | 0.97 (0.81-1) | 0.49 (0.33-0.65) | <0.001 | 0.86 (0.6-0.95) | 0.66 (0.48-0.81) | 0.04 |
| **F1 Score** | 0.99 | 0.83 | <0.001 | 0.88 | 0.57 | <0.001 | 0.95 | 0.63 | <0.001 | 0.82 | 0.79 | 0.002 |

|  | Itching |  |  | Cramp |  |  | Pain |  |  |
| --- | --- | --- | --- | --- | --- | --- | --- | --- | --- |
|  | NLP (95% CI) | ICD (95% CI) | P | NLP (95% CI) | ICD (95% CI) | P | NLP (95% CI) | ICD (95% CI) | P |
| **Sensitivity** | 0.98 (0.86-1) | 0.24 (0.09-0.45) | <0.001 | 0.98 (0.85-1) | 0.09 (0.01-0.29) | <0.001 | 0.99 (0.93-1) | 0.52 (0.37-0.66) | <0.001 |
| **Specificity** | 0.96 (0.8-1) | 0.98 (0.86-1) | 0.68 | 0.89 (0.72-0.98) | 0.98 (0.88-1) | 0.18 | 0.5 (0.93-1) | 0.5 (0.37-0.66) | 1 |
| **PPV** | 0.96 (0.8-1) | 0.8 (0.54-1) | 0.32 | 0.88 (0.69-0.97) | 0.8 (0.16-1) | 0.22 | 0.99 (0.93-1) | 0.98 (0.87-1) | NA |
| **NPV** | 0.98 (0.85-1) | 0.57 (0.41-0.72) | <0.001 | 0.98 (0.86-1) | 0.58 (0.43-0.72) | <0.001 | 0.5 | 0.02 | NA |
| **F1 Score** | 0.97 | 0.56 | <0.001 | 0.91 | 0.28 | <0.001 | 0.99 | 0.68 | <0.001 |

B)
|  | Fatigue |  |  | Nausea and/or vomiting |  |  | Anxiety |  |  | Depression |  |  |
| --- | --- | --- | --- | --- | --- | --- | --- | --- | --- | --- | --- | --- |
|  | NLP (95% CI) | ICD (95% CI) | P | NLP (95% CI) | ICD (95% CI) | P | NLP (95% CI) | ICD (95% CI) | P | NLP (95% CI) | ICD (95% CI) | P |
| **Sensitivity** | 0.98 (0.88-1) | 0.02 | <0.001 | 0.94 (0.79-0.99) | 0.06 (0.008-0.21) | <0.001 | 0.96 (0.8-1) | 0.08 (0.01-0.26) | <0.001 | 0.82 (0.57-0.96) | 0.53 (0.28-0.77) | 0.1 |
| **Specificity** | 0.75 (0.51-0.9) | 0.98 (0.83-1) | 0.6 | 0.95 (0.74-1) | 0.97 (0.82-1) | 0.7 | 0.96 (0.8-1) | 0.96 (0.8-1) | 1 | 0.67 (0.48-0.82) | 0.99 (0.89-1) | 0.002 |
| **PPV** | 0.86 (0.7-0.95) | 0.5 | NA | 0.97 (0.83-1) | 0.8 (0.16-1) | 0.4 | 0.96 (0.8-1) | 0.67 (0.09-0.99) | 0.35 | 0.56 (0.35-0.76) | 0.95 (0.66-1) | <0.001 |
| **NPV** | 0.97 (0.78-1) | 0.4 (0.26-0.55) | NA | 0.9 (0.68-0.99) | 0.4 (0.26-0.55) | <0.001 | 0.96 (0.8-1) | 0.51 (0.36-0.66) | <0.001 | 0.88 (0.69-0.97) | 0.8 (0.65-0.91) | 0.25 |
| **F1 Score** | 0.92 | 0.03 | <0.001 | 0.95 | 0.12 | <0.001 | 0.96 | 0.14 | <0.001 | 0.67 | 0.68 | <0.001 |

|  | Itching |  |  | Cramp |  |  | Pain |  |  |
| --- | --- | --- | --- | --- | --- | --- | --- | --- | --- |
|  | NLP (95% CI) | ICD (95% CI) | P | NLP (95% CI) | ICD (95% CI) | P | NLP (95% CI) | ICD (95% CI) | P |
| **Sensitivity** | 0.91 (0.48-1) | 0.2 (0.005-0.82) | 0.1 | 0.8 (0.16-1) | 0.2* | 0.3 | 0.99 (0.82-1) | 0.13 (0.05-0.26) | <0.001 |
| **Specificity** | 0.93 (0.82-0.99) | 0.99 (0.92-1) | 0.2 | 0.92 (0.8-0.98) | 0.99 (0.93-1) | 0.1 | 0.33 (0.008-0.91) | 0.86 (0.29-1) | 0.3 |
| **PPV** | 0.63 (0.25-0.91) | 0.67 (0.03-1) | 0.26 | 0.33 (0.04-0.78) | 0.5* | NA | 0.96 (0.86-1) | 0.92 (0.54-1) | 0.19 |
| **NPV** | 0.99 (0.92-1) | 0.92 (0.8-0.98) | 0.04 | 0.99 (0.92-1) | 0.96 (0.86-1) | NA | 0.6 (0.03-1) | 0.07 (0.01-0.19) | 0.3 |
| **F1 Score** | 0.74 | 0.31 | 0.008 | 0.47 | 0.29 | 0.03 | 0.97 | 0.22 | <0.001 |

\*Denotes 95% confidence intervals that could not be calculated due to insufficient numbers in cells of the 2x2 tables.

However, specificity was highly variable. In the Bio*Me* dataset, specificity for NLP ranted from 0.5 to 0.96, while specificity for ICD ranged from 0.5 to 0.98. In the MIMIC-III dataset, specificity for NLP ranged from 0.33 to 0.96, while for ICD it was 0.86-0.99. ICD codes were more specific for nausea and/or vomiting (NLP 0.57 vs. ICD 0.97, P=0.03) in BioMe and more specific for depression (NLP 0.67 vs. ICD 0.99, P=0.002) in MIMIC-III. **(Table 2 A/B)**

Twenty-five patients were identified by NLP to have undergone PHQ-9 depression screening. While 3 patients had PHQ-9 scores <10, 2 of the 3 had a clinical history of depression (active group therapy, inpatient psychiatric admissions for depression, or prior suicide attempts) but were receiving adequate treatments therefore resulting in low PHQ-9 scores. The last patient did not have evidence (ICD, NLP, or on chart review) of having depression. Of the 24 patients who were depression positive by PHQ-9 and/or clinical history, NLP correctly identified 22 (92%) patients while ICD 9/10 identified 20 (83%) patients.

### Symptom Burden

In the BioMe cohort, symptom burden was high among HD patients. NLP, identified at least 1 symptoms in 96% of patients, and 4 or more symptoms in 50% of patients. (**Figure 2A)** The number of symptoms identified increased with the number of encounters in BioMe. Patients who did not have any symptoms identified by NLP had a median of 7 encounters (IQR 1-32), while patients with all 7 symptoms had a median of 230 encounters (IQR 141-419). **(Figure 3)** In MIMIC-III, NLP identified at least 1 symptom in 97% of patients, and 4 or more symptoms in 48% of patients. **(Figure 2B)** Encounter analysis could not be performed for MIMIC-III.

**Figure 2:**
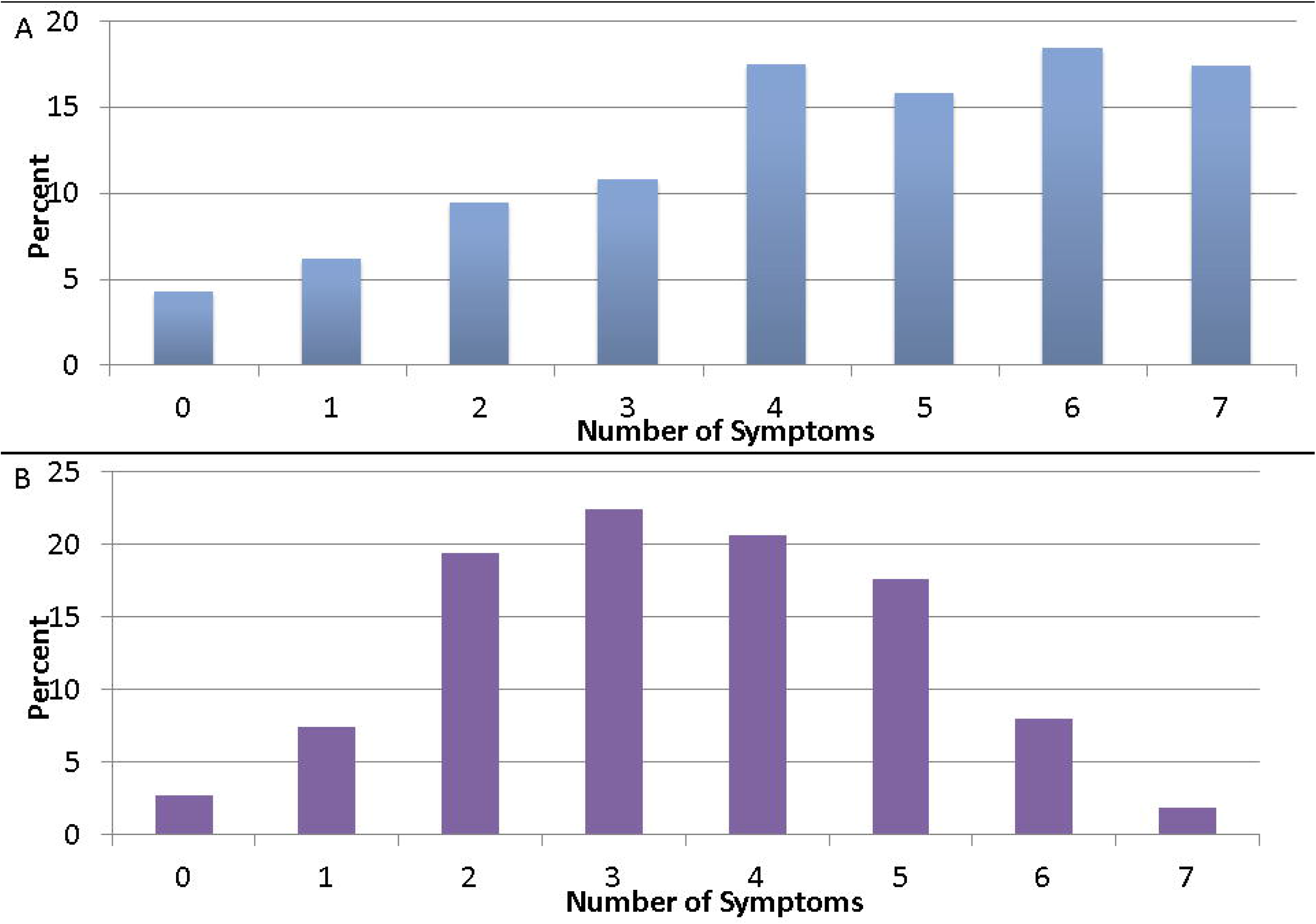
Overall symptom burden of symptoms identified by NLP from (A) Bio*ME* and (B) MIMIC-III.

**Figure 3:**
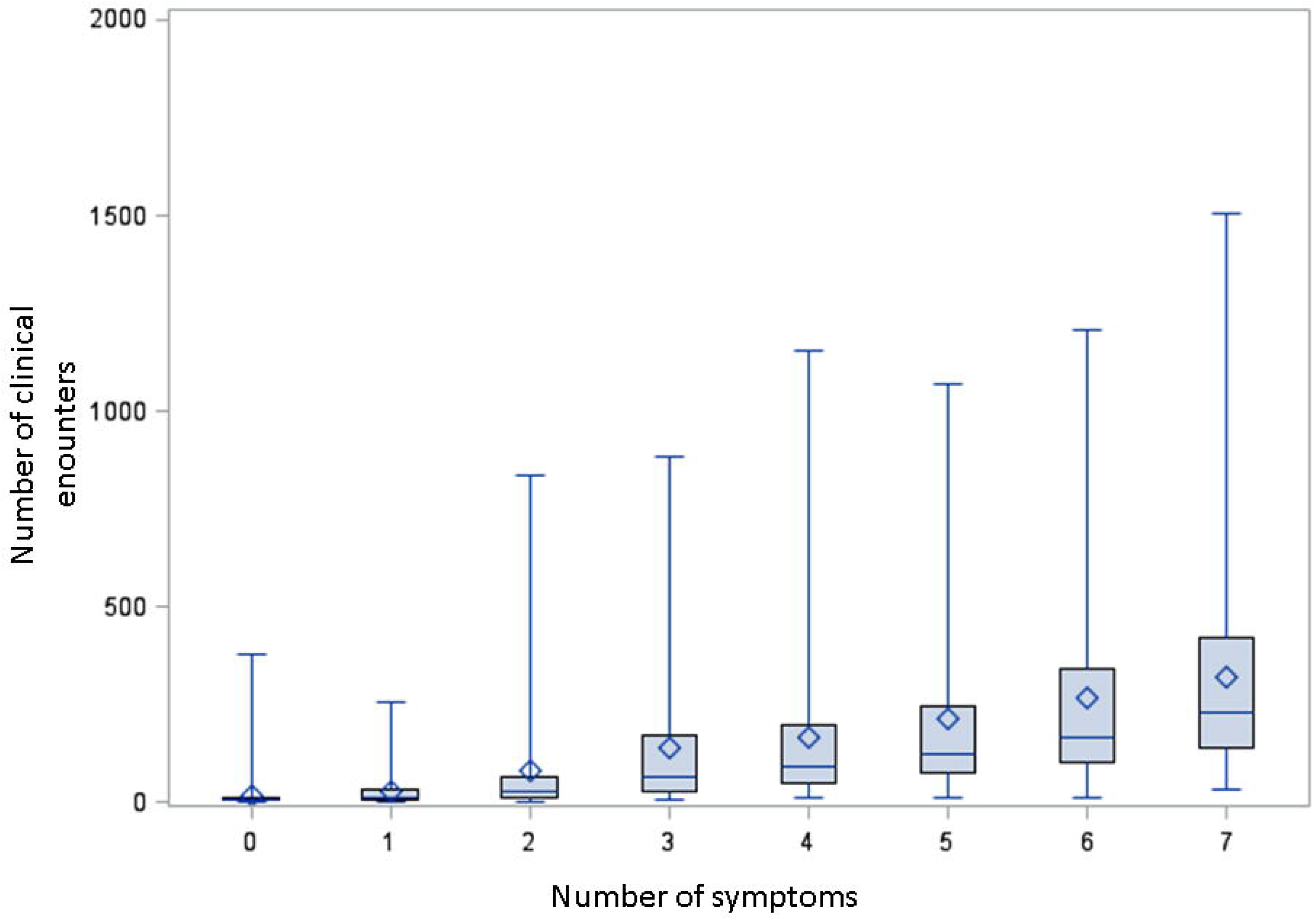
Boxplot demonstrating the number of symptoms identified with the number of clinical encounters.

## Discussion

High-throughput retrospective assessment of symptoms in patients on HD from EHR is difficult. NLP is one potential solution to this problem. We demonstrated that NLP had better sensitivity than ICD codes at identifying seven symptoms with validation across two different cohorts.^3^ The symptom burden was high, with a majority of patients having at least 4 or more symptoms. Finally, identification of symptoms was highly dependent on the number of encounters that HD patients had.

As the care of HD patients is improving, focus has shifted to improving how patients feel (i.e., patient centered outcomes). The SONG-HD initiative has identified several key outcomes important to all stakeholders (patients and physicians) and has emphasized the importance of clinical research that includes these symptoms as both predictors and outcomes. Prior research that employed patient-centered outcomes as endpoints have required prospective surveys for their execution.^2,16^ This is labor-intensive and only allows assessment of symptom burden at the time of the survey. By using NLP, notes can be processed in a high-throughput manner. In addition, benchmarking and reporting these patient-centered outcomes from multiple dialysis providers could provide a unique opportunity to improve clinical practice.

There are currently few studies in nephrology that have utilized NLP. The predominant use has been on the identification of risk factors for progression of chronic kidney disease and the identification of CKD from EHR.^17–22^. Additionally, the studies that have utilized NLP to identify risk factors for progression of chronic kidney disease included few symptoms or patient-centered outcomes in their models. Prior studies in other chronic diseases, such as pancreatic cancer, have demonstrated the utility of NLP to identify the patient-centered outcomes of urinary incontinence and erectile dysfunction from EHR.^23^ Our study supports the use of NLP for identification of patient-centered outcomes in HD patients given the higher sensitivity of the NLP method compared to identification with ICD codes.

We found that overall symptom prevalence identified in the Bio*Me* cohort by NLP is similar to prior published survey data on symptoms, i.e. prevalence of fatigue was reported to be 69-87% in literature and we found a prevalence of 84%.^16,24,25^ Certain symptoms were less commonly found such as itching (NLP 48% vs. literature 52-70%) and cramps (NLP 45% vs. literature 43-74%), while other symptoms such as nausea and/or vomiting (NLP 74% vs literature 26-35%) were more commonly found. The differences are likely due to differences in cohorts and settings.

While surveys are done in patients who are stable at their outpatient hemodialysis centers, BioMe provider notes consist not only of preventative care visits, but also acute inpatient and outpatient notes where more severe symptoms are documented. As MIMIC-III consists of progress notes from hospital visits that required an ICU admission, symptoms were identified at an even lower rate. One potential reason is that patients admitted to ICUs are critically ill and potentially with altered mental status or mechanically ventilated which prevents patients from verbalizing their symptoms. Additionally, patient care and billing is often focused on the admission diagnosis and contributing comorbidities, while symptoms and psychosocial comorbidities may not be as well addressed in notes. We chose to not to place limitations on the number, timing, or type of notes, which may have increased the likelihood of NLP identifying a symptom. However, comparator measures via ICD 9/10 codes, were also identified without limitations to encounters, allowing for a fair comparison.

Despite these differences in cohorts, NLP was significantly more sensitive than ICD codes for identification of nearly all symptoms in both Bio*Me* and MIMIC-III. ICD9/10 codes are commonly used for the identification of several disease processes from administrative data and we found that NLP out performed ICD 9/10 codes at identification of all symptoms in both BioMe and MIMIC-III.^26–29^ As ICD 9/10 codes are administrative codes clinicians may be less inclined to use them to document symptoms experienced by HD patients. When ICD codes for symptoms were present in our data, they identified symptoms with high specificity.

While NLP was more sensitive at identifying depression, ICD codes were more specific. A substantial portion of the false positives for depression was due to the use of depression in other clinical contexts. As there was no consistent way that this was documented across notes it could not be easily addressed in our NLP algorithm.

Our study should be interpreted in light of some limitations of our study including the dependence of symptom identification on the number of encounters and notes available, the more encounters available the more likely a provider was to document a symptom. However, this is a common issue with EHR systems, where both sicker patients as well as patients with longer length of follow up having more data.^30^ Additionally, only symptoms which the provider are screening for are documented and therefore NLP may miss those symptoms patients are not discussing with their providers. Neither the BioMe nor MIMIC-III datasets are exclusive to outpatient HD patients, which make comparison with prior published data on outpatient HD patients difficult. However, the prevalence of symptoms in our study is similar to prior published survey data.^24,25^ Unfortunately, as we did not have concurrent survey data available, we used manual chart review as our gold standard. We did extract PHQ-9 survey results to further validate our findings, however only a small portion of patients had this screening done. Additionally, the results of our sensitivity, specificity, PPV, NPV, and F1 scores were relatively consistent across the BioMe and MIMIC-III cohort, suggesting that our NLP algorithm would have good generalizability across different medical systems.

In conclusion, we utilized NLP to identify important patient symptoms from EHR of HD patients from two diverse medical systems. Prevalence of symptoms identified by NLP was similar to previously published survey studies. NLP out performed ICD codes for identification in regards to sensitivity, NPV, and F1 score for a majority of symptoms in both cohorts. Additional refinement of the NLP algorithm and testing in the EHR of outpatient HD units is needed to further validate our findings.

Author Contributions: LC, SC, and GNN designed the study. TVV parsed the data. LC, KC, and ND carried out the analysis. LC and KB performed the manual chart review. ND and AS made the figures and tables. All authors drafted and revised the manuscript and approved the final version of the manuscript.

## Acknowledgements

We want to thank all participants of Bio*Me* and MIMIC-III.

